# Wastewater metaproteomics: tracking microbial and human protein biomarkers

**DOI:** 10.1101/2025.02.08.637285

**Authors:** Claudia G. Tugui, Filine Cordesius, Willem van Holthe, Mark C.M. van Loosdrecht, Martin Pabst

**Affiliations:** Department of Biotechnology, Delft University of Technology, 2629 HZ Delft, The Netherlands

**Keywords:** metaproteomics, wastewater, wastewater-based epidemiology, biomarkers, gut microbes

## Abstract

Wastewater-based surveillance has become a powerful tool for monitoring the spread of pathogens, antibiotic resistance genes, and measuring population-level exposure to pharmaceuticals and chemicals. While surveillance methods commonly target small molecules, DNA, or RNA, wastewater also contains a vast spectrum of proteins. However, despite recent advances in environmental proteomics, large-scale monitoring of protein biomarkers in wastewater is still far from routine. Analyzing raw wastewater presents a challenge due to its heterogeneous mixture of organic and inorganic substances, microorganisms, cellular debris, and various chemical pollutants. To overcome these obstacles, we developed a wastewater metaproteomics approach including efficient protein extraction and an optimized data-processing pipeline. The pipeline utilizes de novo sequencing to customize large public sequence databases to enable comprehensive metaproteomic coverage. Using this approach, we analyzed wastewater samples collected over three months from two urban locations. This revealed a core microbiome comprising a broad spectrum of microbes, gut bacteria and potential opportunistic pathogens. Additionally, we identified nearly 200 human proteins, including promising population-level health indicators, such as immunoglobulins, uromodulin, and cancer-associated proteins.

## Introduction

Globally, approximately 380 trillion liters of wastewater are produced annually, and with the steadily growing world population, it is estimated to nearly double in the next 50 years^1^. Wastewater streams are a complex collection of chemicals, organic compounds, microorganisms, and biomolecules such as DNA and proteins, of which a large fraction originates from human activity. The analysis of wastewater for microbial pathogens, viruses, and substances such as pharmaceuticals, pesticides, and biomarkers of stress and diet has become a routine practice. This has been termed wastewater-based epidemiology (WBE) by Cristian G. Daughton in 2001^2-4^. Today, WBE includes various biological biomarkers, to assess the health status at a population level^5^. Wastewater-based epidemiology (WBE) has proven to be effective for identifying and monitoring epidemic outbreaks. For example, in the 1980s, wastewater surveillance in Finland and Israel provided insights into the spread of the poliovirus^6 7^. Furthermore, during the Coronavirus pandemic, various research groups and governments established COVID-19 surveillance programs^8 9 10^. This informed governmental bodies and the general public about the spread of SARS-CoV-2^11, 12^. Furthermore, the presence of certain bacteria also informs on the spread of antimicrobial resistance, and various diseases^13-17 18 19^.

Apart from the advantage of anonymity, the collection of wastewaters is relatively cheap, and it can be applicable to a large population size. The detection of small molecules such as pharmaceuticals employs chromatographic separation combined with mass spectrometry^20^. The analysis of viruses, microbes or antimicrobial resistance genes commonly employs targeted approaches such as various nucleic acid–based polymerase chain reaction methods^21-26^. Recently, untargeted methods using next-generation sequencing methods have become more affordable and increasingly popular for studying water and wastewater environments^24, 27-30^.

In addition to small molecules, microbes and viruses, wastewater also contains excreted human proteins and proteins from food waste or agricultural activities. Interestingly, many biomarkers that potentially contain information about population health are proteins, excreted through saliva, stool or urine. Currently, a transition toward precision medicine is underway, which prioritizes proactive, patient-centered approaches^31, 32^. One objective is to identify protein biomarkers that can improve early diagnosis^32^. For example, the proteins found in urine can indicate urogenital disorders, chronic conditions like cancer^33 34 35^, autoimmune diseases^36^, neurological disfunctions like Alzheimer^37^ as well as diabetes^38^. However, currently, large volumes of clinical data from biofluids, such as urine and blood, must be collected and analyzed, ideally with minimal discomfort for patients^32^. Wastewater, on the other hand, is readily available and can be used by health professionals to assess the overall health of the population in a simple, non-invasive, and anonymous manner.

While the analysis of small molecules and the targeting of RNA and DNA have become routine, effective protocols for large-scale monitoring of macromolecules, such as proteins, are still lacking. Over the past decades, mass spectrometry-based proteomics has evolved from focusing on single species to encompassing the field of microbial ecology, known as metaproteomics. Metaproteomics enables the measurement of complex microbial mixtures, providing insights into the microbial composition and expressed microbial functions^39-41^. Furthermore, it allows to measure freely floating proteins, including those excreted by humans or released through industrial and agricultural activities. Therefore, metaproteomics can provide an alternative view on wastewater, which cannot be obtained by DNA-based approaches alone. Recent advancements in mass spectrometric instrumentation have significantly reduced measurement time, even for highly complex metaproteomic samples, while also enhancing sensitivity^42, 43^. This is a significant step toward establishing metaproteomics as a routine, untargeted wastewater surveillance approach. However, the heterogeneous nature of wastewater presents additional challenges. First, an effective sample preparation method is required to capture all proteins present. Second, data processing requires a reference sequence database that includes all proteins in the wastewater. While whole metagenome sequencing covers the microbial population, it does not capture freely floating proteins or those from food waste residues and agricultural activities^44, 45^. Additionally, although increasingly affordable, whole metagenome sequencing is time-consuming and prone to errors at various stages, including DNA extraction and data processing^45, 46^.The first metaproteomic study on wastewater, to the best of the authors’ knowledge, was conducted by Carrascal and co-workers who used polymeric adsorbents immersed in the influent water of a wastewater treatment plant over several days^47^. The sorbed proteins allowed for the identification of 690 proteins from bacteria, plants, animals, and humans. In addition to the polymeric probe, the study utilized a large, generic database for database searching. This was later combined with the regions of interest multivariate curve resolution approach, to streamline data analysis^48^. Subsequent studies performed separate analysis of soluble and particulate fractions and concentrating of larger volumes of wastewater followed by SDS-PAGE gel electrophoresis and in-gel digestion to characterize the wastewater proteome^49, 50^. These studies identified various proteins, including potential human biomarkers, as well as a spectrum from various microbes. The proteomic profiles also provided insights into the presence of local industries, such as farming. However, polymeric probes may not capture all freely floating proteins, and separating fractions and concentrating large volumes of wastewater can be time-consuming. Additionally, the choice of reference sequence database affects the accuracy and comprehensiveness of the results. Using generic databases is computationally intensive and may reduce the sensitivity of the database search approach.

In this study, we demonstrate a streamlined metaproteomics approach which we applied to crude municipal wastewater samples collected over three months from two different locations. We developed an efficient sample preparation procedure that extracts proteins from both insoluble and soluble fractions starting with small volumes of wastewater, making it suitable for multiplexing. Additionally, we created a wastewater metaproteomics data processing pipeline that employs *de novo* sequencing to focus generic reference sequence databases in order to obtain a comprehensive metaproteomic coverage.

## Material and Methods

**Sampling**. Samples were taken from two wastewater treatment plants over a period of 4 months, from November 2023 to February 2024. From the wastewater treatment plant Harnaschpolder^6^ samples were taken on 29/11/23, 12/05/23, 24/01/24, 31/01/24, and 20/02/24, and from Utrecht (UT), samples were taken on 29/11/23, 05/12/23, 25/01/24, 23/02/24 and 27/02/24. The sampling was done from the influent, raw sewage, on the days with low precipitation. After sampling influent wastewater, samples were stored at -20 ^°^C until further processed by a short sample preparation protocol. **Protein extraction and proteolytic digestion**. 500 µL of the wastewater influent was taken and diluted with B-PER (175µl) and 50 mM TEAB buffer (175 µL) and heated at 90 °C, for 5 min under shaking at 300 rpm. Further, the sample was subjected to cell lysis using vortexing 3 times for 1 minute using a bench vortexing machine, sonication on a sonication bath for 15 minutes and one freeze/thaw cycle (frozen at -80 °C, thawed in incubator at 40°C for 5 minutes). The samples were then centrifuged and transferred to a 1.5 mL LoBind Eppendorf tube. TCA was added to the sample at a ratio 1:4 (v/v, TCA/sample), vortexed and incubated at 4 °C for 20 minutes. After centrifugation, the protein pellet was re-solubilized in 6 M urea and then reduced with DTT (dithiothreitol) and alkylated using IAA (iodoacetamide). After alkylation, the sample was transferred to a FASP filter (Millipore, MRCPRT010) which was previously conditioned by washing 2 times with 100 mM ABC buffer. The filters were centrifuged at 14K rpm in a bench top centrifuge, for 45 minutes, and then 2 times at 14K rpm for 40 minutes after adding 100 mM ABC buffer. Next, the proteins were proteolytically digested on the FASP filter by adding 100 µL trypsin solution, which was prepared by diluting 8 µL trypsin stock solution (0.1 ug/mL in 1 mM HCl, Promega, Cat No) in 100 µL 100 mM ABC. The FASP filters were incubated over night for digestion, at 37 °C, under gentle shaking at 300 rpm. The following day, the filters were centrifuged and then once washed with 100 mM ABC buffer followed by a second wash with 100 µL of 10% ACN 0.1% FA/H_2_O collected the proteolytic peptides. The pooled fraction was then purified using an OASIS HLB well plate (Waters, UK) according to the manufacturer’s protocol. The purified peptide fraction was speed-vac dried and stored at -20 °C until further analyzed. **Shotgun metaproteomics**. To the speed vac tried samples 20 µL of 3% acetonitrile and 0.01% trifluoroacetic acid in H_2_O was added, vortexed, then left at room temperature for 30 min, and then once more vortexed. The peptide concentration was determined by measuring the absorbance at 280 nm using a NanoDrop ND-1000 spectrophotometer (Thermo Scientific). Samples were diluted to a concentration of approximately 0.5 µg/µL. Shotgun metaproteomics was performed as described previously^41^, with a randomized sample order. Briefly, approximately 0.5 µg protein digest was analysed using a nano-liquid-chromatography system consisting of an EASY nano-LC 1200, equipped with an Acclaim PepMap RSLC RP C18 separation column (50 μm x 150 mm, 2 μm, Cat. No. 164568), and a QE plus Orbitrap mass spectrometer (Thermo Fisher Scientific). The flow rate was maintained at 350 nL/min over a linear gradient from 5% to 25% solvent B over 90 min, from 25% to 55% over 60 min, followed by back equilibration to starting conditions. Solvent A was a 0.1% formic acid solution in water (FA), and solvent B consisted of 80% ACN in water and 0.1% FA. The Orbitrap was operated in data dependent acquisition (DDA) mode acquiring peptide signals from 385–1250 m/z at 70 K resolution in full MS mode with a maximum ion injection time (IT) of 75 ms and an automatic gain control (AGC) target of 3E6. The top 10 precursors were selected for MS/MS analysis and subjected to fragmentation using higher-energy collisional dissociation (HCD) at a normalised collision energy of 28. MS/MS scans were acquired at 17.5 K resolution with AGC target of 2E5 and IT of 75 ms, 1.2 m/z isolation width. **Taxonomic profiling and database construction**. The mass spectrometric raw data for each sample were de novo sequenced using PEAKS Studio X (Bioinformatics Solutions, Inc., Canada). De novo sequences with an ALC score >70 were subjected to taxonomic profiling using the NovoBridge pipeline as described previsously^51^. An in-house constructed API sequence downloader “UniRefBuilder” was employed to construct a reference sequence database containing all UniRef90 entries of the identified families per sample. The NovoBridge+ pipeline and the UniRefBuilder are freely available via GitHub: https://github.com/hbckleikamp/NovoBridge_plus, and https://github.com/claudiatugui/UniRefBuilder. **Database searching**. The focused UniRef90 database was used for database searching using PEAKS Studio X (Bioinformatics Solutions, Inc., Canada) employing a two-round search approach. The first round allowed for one missed cleavage and included carbamidomethylation as a fixed modification, allowing a 20 ppm precursor error and a 0.02 Da fragment ion error. From every sample, the matched proteins from the first round search (without score cut-offs) and the proteins from the human reference proteome (UP000005640) were combined into a new reference sequence database for the second-round search. The second-round search was performed allowing up to 3 missed cleavages, with carbamidomethylation as a fixed modification, and methionine oxidation and asparagine or glutamine deamidation as variable modifications, allowing 20 ppm precursor error and 0.02 Da fragment ion error. Peptide-spectrum matches from the second round were filtered to a 5% false discovery rate (FDR) at the PSM level, and protein identifications with ≥2 unique peptide sequences were considered significant. The complete dataset was combined using the PEAKSQ module allowing 10 minutes RT shifts and 10 ppm mass error. All identified proteins were exported and further processed as described in the following. **Data analysis and visualization**. For further analysis, proteins were filtered for a minimum of 2 unique peptides and an overall top 3 peptide area (summed across all samples) of > 5E5, and finally only the top hit from every protein group was kept for further data visualization and interpretation. The protein identification table was further analyzed in Python. A taxonomic lineage based on NCBI taxonomy was assigned to every protein which was then used to prepare a taxonomic composition at different taxonomic levels. Two datasets were generated, the “microbial dataset” containing all proteins excluding those with “Eukaryota” and missing annotations at the superkingodm level, and the “human proteome” dataset which contains only protein identifications with the annotation “Homo” at the genus level. Finally, for both datasets, only one protein per protein group was retained (“one_per_group” datasets). Except stated otherwise, the taxonomic composition, protein abundance and diversity plots were determined by summing the top 3 peptide areas for each protein within the respective taxonomy. Principal coordinate analysis (PCoA) was performed using the MDS implementation from scikit-learn, after computing the Bray-Curtis dissimilarity matrix based on genus-level compositions from all time points and locations. The Shannon diversity index was calculated in Python using the formulae H′=−∑piln(pi), where pi is the proportional abundance of each genus. The microbial proteins were categorized according to human gut bacteria and potential pathogens. Assigning potential pathogenic bacterial genera was based on the work by Bartlett et al.^52^. The assignment of human gut microbes was performed using the Human Gut Microbiome Atlas (www.microbiomeatlas.org) which was queried via an API. Annotation of potential pathogenic microbes was done according to the WHO report^53^. The human dataset was further analyzed for the enrichment of molecular and cellular functions, and pathways using STRING^54^. Potential cancer related protein biomarkers were taken from the Human Protein Atlas (www.proteinatlas.org)^55^, which was queried via an API. Human proteins associated with coronary artery disease (CAD) were taken from the CAD biomarkers database^56^, diabetic nephropathy (DN) from Zürbig *et al*. (2012)^57^, breast cancer from Beretov *et al*. (2015)^58^, urothelial cancer from Chen *et al*. (2021)^59^, Abdominal-type Henoch-Schonlein purpura (HSP) from Jia *et al*. (2021)^60^, prostate cancer from Fujita *et al*. (2018)^61^, and for ovarian cancer and IBD from Owens *et al*. (2022)^62^. Parts of Figure 1A were generated with BioRender (Created in BioRender. Tugui, C. (2025) https://BioRender.com/v22q517). All mass spectrometric proteomics raw data are available via the ProteomeXchange consortium database, through the identifier PXD059455.

**Figure 1.**
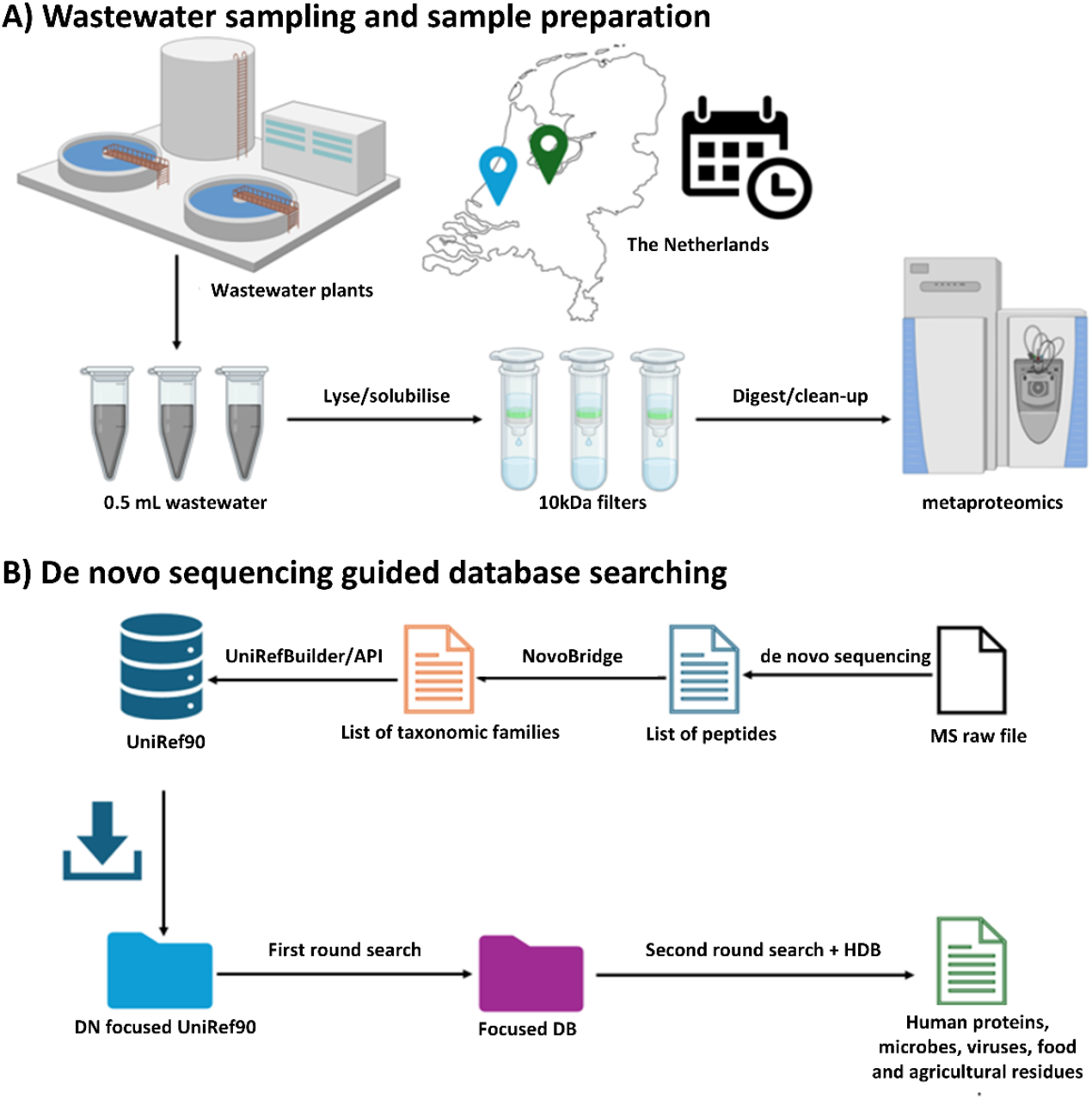
The wastewater metaproteomics workflow involved rapid protein extraction and de novo sequencing guided database searching to identify a broad spectrum of proteins across all domains of life. Influent samples were collected from two wastewater treatment plants, located in Utrecht and Harnaschpolder (The Netherlands), over a 3-month period, with five time points sampled from each plant. A 0.5 mL aliquot of influent was subjected to cell lysis, protein extraction, and proteolytic digestion using a filter- aided sample preparation (FASP) method. Shotgun metaproteomics was performed using a hybrid quadrupole-Orbitrap mass spectrometer. The resulting data were de novo sequenced to focus the global UniRef90 database. This enabled a rapid two-step database searching using the global Uniref90 database content to enhance protein identification.

## Results

Wastewater samples from 2 different wastewater treatment plants were sampled during the winter months over a time period of 3 months (Dec 2023 – Feb 2024). The wastewater treatment plant in Harnaschpolder is the biggest wastewater treatment plant in The Netherlands, serving over 1 million inhabitants and 40.000 companies in The Hague area. The wastewater treatment plant located very close to the city of Utrecht serves the city and the surrounding area of around 375.000 people. The main difference between both sampling locations, apart from the size, is their proximity to an urban area. The wastewater treatment plant located in Utrecht is close to the city serving a highly urbanized area, with relatively low sewer residence times. Harnaschpolder, on the other hand, is located between The Hague and Delft, in a peri-urban zone including different larger and smaller industries. The wastewater is transported over longer distances by pressure mains. To capture a comprehensive spectrum of taxonomies from the sampled wastewater, we developed a streamlined wastewater metaproteomics workflow. This included the extraction and analysis of proteins from both the liquid and solid components from small volumes of influent wastewater (raw sewage). For this, 0.5 mL of wastewater was directly mixed with a lysis buffer, followed by solubilization and cell lysis using a combination of sonication and freeze-thaw cycles. Proteins were then extracted and digested using a filter-aided sample preparation (FASP) workflow.

The proteolytic peptides were finally analyzed with a shotgun metaproteomics experiment employing a one-dimensional chromatographic separation (with a 120 minutes gradient) and a hybrid quadrupole-Orbitrap mass spectrometer. This allowed to detect 4,919 proteins across 3,009 protein groups, including 249 human proteins from 198 protein groups, when considering proteins with ≥2 unique peptides. The number of proteins increased to >10,000 when considering proteins with only 1 unique peptide. The identified proteins could be assigned to a diverse array of taxonomies and sources, including environmental microbes, human microbiome microbes (both pathogenic and commensal bacteria), human and animal proteins, as well as agricultural waste and food residues.

## Wastewater metaproteome biomass composition and diversity

The most abundant proteins in the wastewater across both treatment plants could be associated with human and animal feces, urine, and other animal-related sources. Although these proteins constituted only a minor fraction of the total protein fraction (approximately 10–15%), these accounted for more than 50% of the total protein abundance (considering the top 3 peptide area per protein) (Figure 2A).

**Figure 2:**
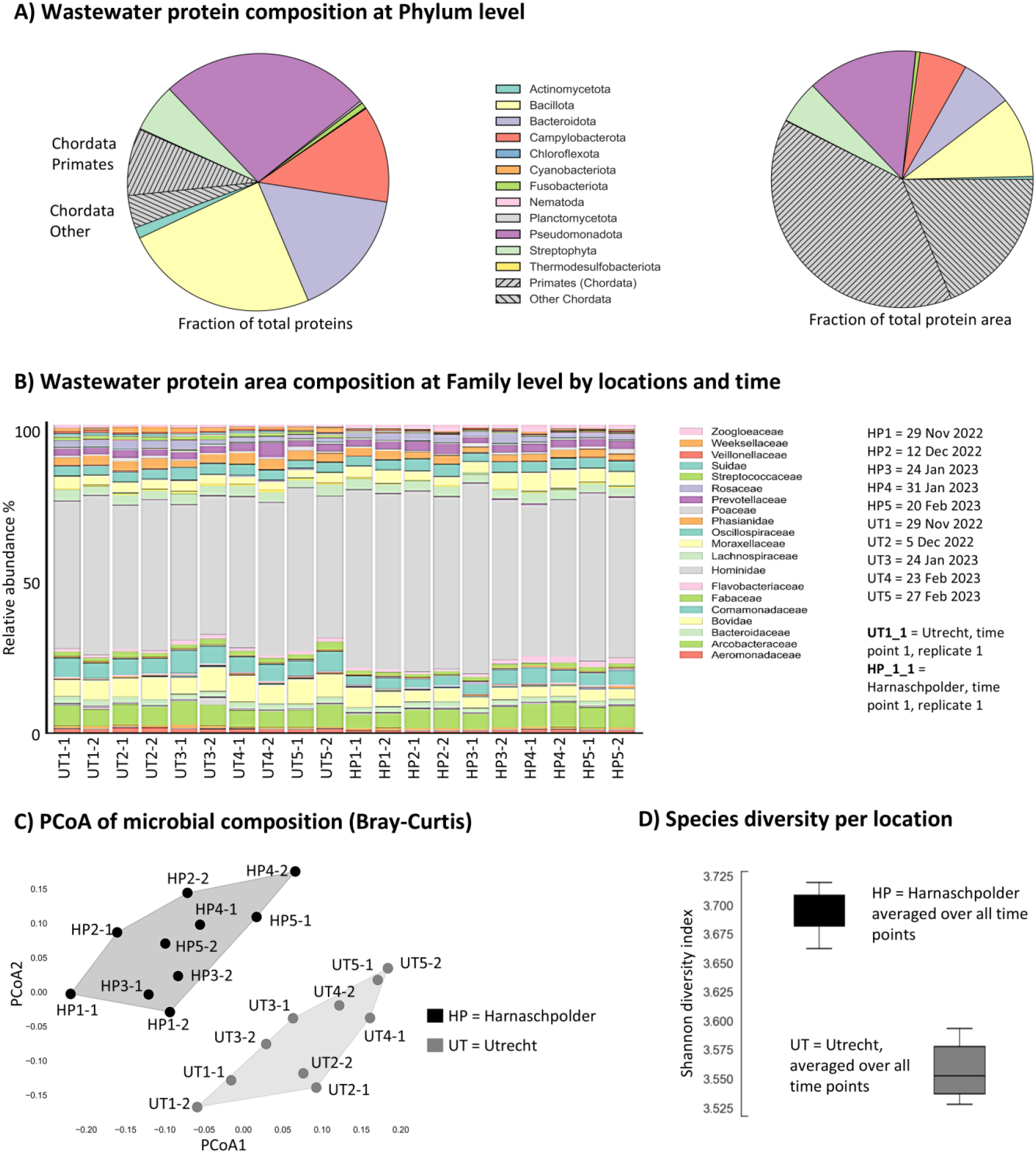
A. The graph displays the averaged phylum-level distribution from both locations, HP and UT. This is represented either by summing the top three peptide areas of all proteins (“Fraction of total protein”) or by counting the number of identified proteins per phylum (“Fraction of total protein area”). B. The bar graph illustrates the protein area composition at the family level across 5 time points from Dec 2023 to February 2024, observed in the wastewater of Harnaschpolder ^6^ and Utrecht (UT), located in The Netherlands. C. The graph depicts alpha diversity, calculated for bacterial genera using the Shannon diversity index. D. The Principal Coordinate Analysis (PCoA) plot visualizes beta diversity, calculated by Bray-Curtis dissimilarity, using all microbial proteins detected in the samples.

Among the dominant microbial phyla, we identified *Pseudomonadota, Bacillota, Bacteroidota*, and *Campylobacterota*, which are typical wastewater-associated microbes or microbes associated with various niches of the human microbiome.

Minor other phyla that were consistently identified included *Actinomycetota, Fusobacteriota, Planctomycetota*, and *Cyanobacteriota*, which likely contribute to organic matter degradation and nutrient cycling in these environments. Additionally, significant amounts of *Streptophyta* were detected which are plant-derived residues, possibly from food processing activities. Finally, we also detected *Nematoda* which is a diverse phylum of worms commonly found in soil, and aquatic environments, and as parasites in plants and animals.

Interestingly, these Phyla could be further assigned to >60 taxonomic families, which indicates a diversity and complex ecosystem. Nevertheless, the individual sampling time points during the winter period, as well as the different locations (HP and UT), showed only comparatively small differences in their core taxonomic profiles (Figure 2B).

The most dominant families derived from fecal contamination such as *Streptococcaceae, Enterobacteriaceae, Bacteroidaceae, Veillonellaceae*, and *Bifidobacteriaceae*, which are indicative of human and animal gut microbiota and include potential pathogens like Streptococcus and *E. coli*^*63*^. These also included opportunistic pathogens, such as *Weeksellaceae, Leptotrichiaceae, Aeromonadaceae*, and *Arcobacteraceae*. Other families such as *Planctomycetaceae, Nitrobacteraceae, Comamonadaceae*, and *Rhodospirillaceae*, are rather related to biological processes including nitrogen cycling and organic matter degradation^64-66^. Additionally, biodegraders like *Pseudomonadaceae, Bacillaceae, Chitinophagaceae, Flavobacteriaceae, Burkholderiaceae*, and *Sphingomonadaceae* were detected which play crucial roles in the breakdown of organic material and xenobiotics^67-72^. Interestingly, also *Caldilineaceae* and *Nocardioidaceae* were detected which are frequently linked to operational challenges in wastewater treatment, such as sludge bulking and foaming^73, 74^. Also, *Zoogloeaceae* were detected *Zoogloeacease* is a major denitrifying bacterium which also produces extracellular polymeric substances relevant for floc formation^75^. Members of the families *Moraxellaceae* and *Burkholderiaceae* have been frequently linked to opportunistic infections^76-81^.

Non-microbial taxa could be assigned to food (processing) residuals, livestock and agricultural runoff, which included families such as *Suidae, Bovidae, Phasianidae*, and *Callorhinchidae*. A broad range of different plant families were also detected including *Arecaceae, Asteraceae, Fabaceae, Solanaceae, Poaceae, Malvaceae, Zingiberaceae, Cucurbitaceae, Rosaceae, Anacardiaceae, Juglandaceae*, and *Pedaliaceae* which likely derive from food residuals, and agricultural activities, and which contribute organic material in wastewater environments. However, also proteins from potential pathogenic vectors such as *Nematoda* (e.g., *Steinernematidae*^82^) were detected, as well as from *Blastocystidae* which is the most prevalent gastrointestinal protist in humans and animals. While its clinical significance remains uncertain, it is increasingly regarded as a commensal component of the gut microbiome^83^. Overall, a large fraction (approx. 50%) of proteins derived from families that include potential pathogens, including several that are linked to the gut microbiome. A comprehensive list of identified proteins, Phyla and Families is provided in the SI EXCEL DOC.

To further investigate the similarity and differences between the individual sampling time points and locations, we performed a principal coordinates analysis (PCoA) based on the Bray-Curtis dissimilarity index (Figure 2C). In general, in PCoA, points that are closer together on the plot represent taxonomic profiles with more similar compositions. The analysis demonstrated a clear clustering of replicate samples, and while the individual sampling time points showed distinct taxonomic profiles, the samples also clearly clustered by location. When further analyzing for differences between both locations, the UT wastewater generally showed a slightly lower alpha-diversity compared to the HP wastewater, with a Shannon index of 3.55 and 3.75, respectively (Figure 2D). The Shannon index is a metric that reflects both species richness (the total number of species) and the evenness of their abundances within a community. Higher values, such as detected for the analyzed wastewater, indicate a more diverse community with a larger number of species and relatively balanced abundances among them. However, considering that metaproteomics requires a minimum of protein biomass for a successful detection of a microbe, the true complexity is likely significantly larger.

We compared the average abundance of selected microbes between both locations, which showed that several bacterial families differed in their relative abundance (Figure 3). For example, in HP, *Aeromonadaceae* and *Weeksellaceae*, which are common in wastewater and are associated with potential pathogenic species^84^ and are known to harbor antibiotic resistance genes or coexist with microbes involved in the spread of antimicrobial resistance^84-86^, showed a much higher relative abundance in HP compared to UT. Additionally, *Streptococcaceae* and *Vibrionaceae* were more abundant in HP, both of which are fecal indicators and potential pathogens^87, 88^. In contrast, *Sphingomonadaceae*, which are well-known biodegraders of xenobiotics in wastewater^70, 72^, were more abundant in the UT samples. On the other hand, *Nocardiaceae* and *Nocardioidaceae* were present at comparable levels in both locations. *Nocardiaceae* are known to cause foaming issues in wastewater treatment plants^89, 90^, while the other includes members, such as *Nocardioides* which can degrade a variety of pollutants^91^. Overall, a significant proportion of the detected microbial families could be associated with pathogens and diseases. However, the unambiguous identification of pathogens requires species-level resolution, which could not be achieved using the generic reference sequence database.

**Figure 3:**
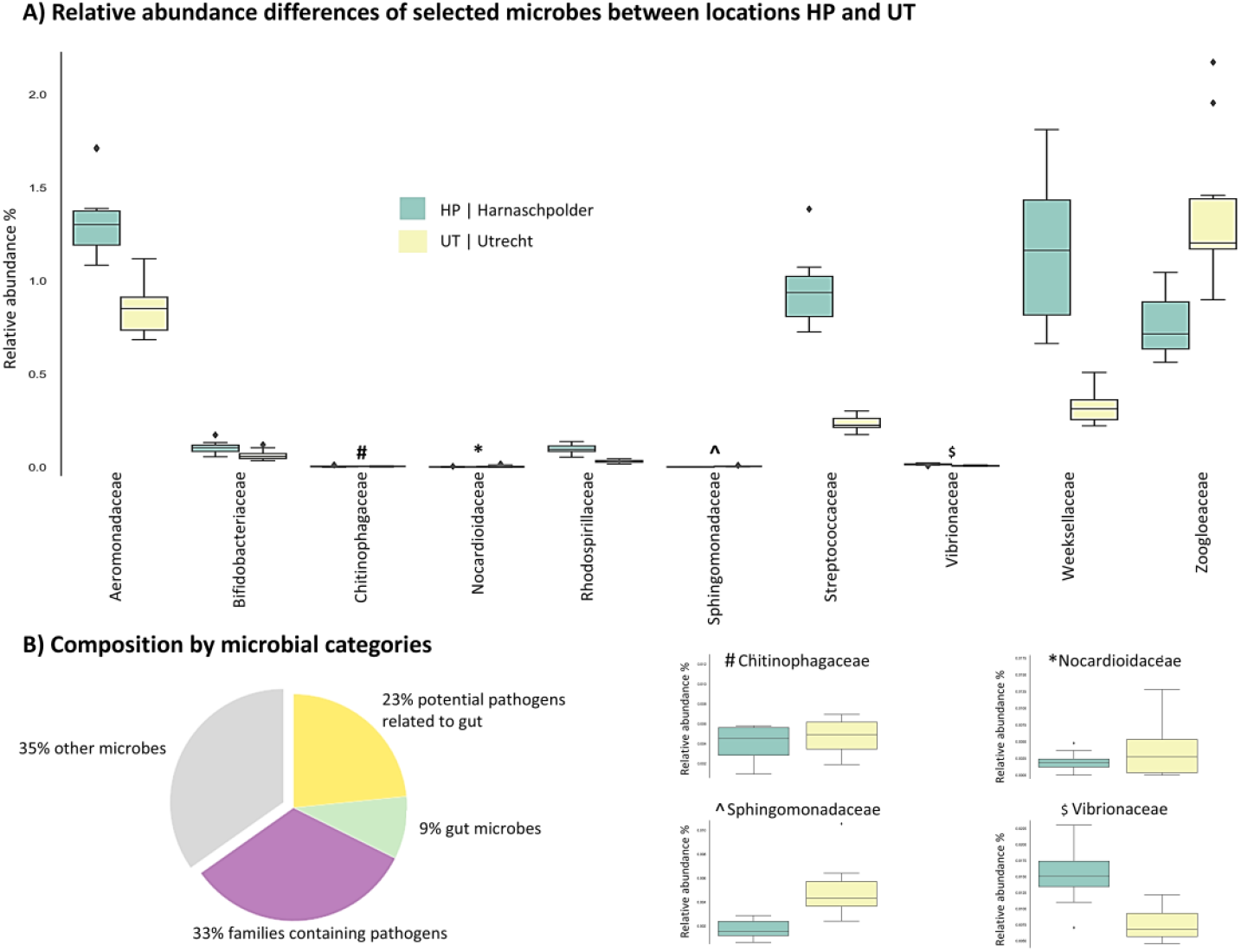
Graph A displays boxplots of the relative abundance of selected bacterial families across 5 time points from Dec 2023 to February 2024, observed in the wastewater of Harnaschpolder^6^ and Utrecht (UT), located in The Netherlands. Graph B presents a pie chart illustrating the distribution of microbes observed in the wastewater, categorized into groups, by: i) those associated with pathogens, ii) gut microbes, iii) both and iv) other microbes.

## The human wastewater proteome

Over 190 proteins groups from the human proteome were detected across all sampling time points and wastewater locations, which could be assigned to sources such as feces, urine, sweat, or saliva (SI EXCEL DOC). Interestingly, the human proteome profile was comparable across all sampling time points and locations. Furthermore, only a subset of these proteins exceeded 2.5% relative abundance, with chymotrypsin-like elastase family member 3A being the most abundant, which accounted for approximately 15% of the total abundance of all detected human proteins. Other abundant proteins did not exceed a relative abundance of 2.5%, including keratin type II cytoskeletal 1, chymotrypsin-C, keratin type I cytoskeletal 9, uromodulin, albumin, alpha-1-antitrypsin, immunoglobulin J chain, keratin type I cytoskeletal 10, and pancreatic alpha-amylase, while most others remained below 1% relative abundance (Figure 4, and SI EXCEL DOC). Nevertheless, many of these proteins are promising biomarkers for accessing the health status or detecting diseases within the general population. Multiple proteins could be associated with breast cancer, intestinal diseases, pancreatic cancer, and gastrointestinal cancers, including stomach carcinoma and colon cancer. Cancer- related associations link to various types of carcinomas, such as adenocarcinoma, breast cancer, pancreatic carcinoma, and skin carcinoma. Additionally, this also includes associations with autoimmune diseases, genetic diseases, and metabolic disorders, including diabetes mellitus. Several diseases related to immune system dysfunction were also enriched. Furthermore, infectious diseases, including bacterial infectious disease and viral infectious disease, were also found, reflecting the role of these proteins in immune responses to infections. The complete tables of enriched terms and functions is provided in the SI EXCEL DOC. A more detailed study regarding proteins related to cancer with annotations from the Human Protein Atlas can be found in the SI DOC (SI Figure 2 and SI Table 1). An overview of potential biomarkers for widespread diseases is provided in Figure 5C, and SI EXCEL DOC, and SI DOC (SI Table 1).

**Figure 4:**
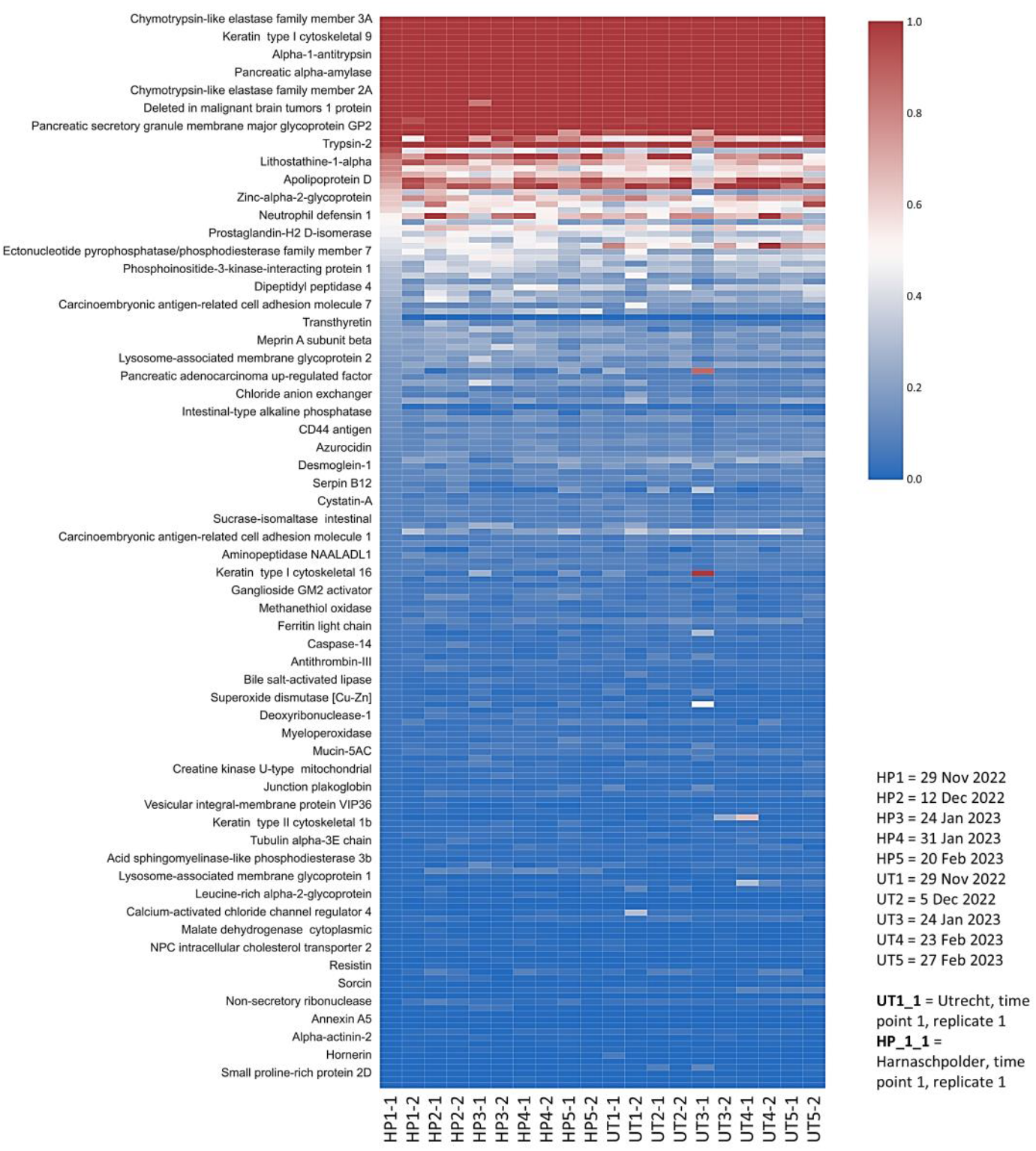
The heatmap provides an abundance profile of all human proteins identified in the wastewater, across 5 time points from Dec 2023 to February 2024, observed in the wastewater of Harnaschpolder^6^ (HP) and Utrecht (UT), located in The Netherlands. The color gradient represents the relative abundance of each human protein, determined by the summed top-three peptide areas. The summed area of all proteins was normalized to 100. The proteins were further sorted by ascending abundance, based on sample HP1-1 (Harnaschpolder, time pointe 1 replicate 1). The y-axis annotation names every fourth protein.

**Figure 5:**
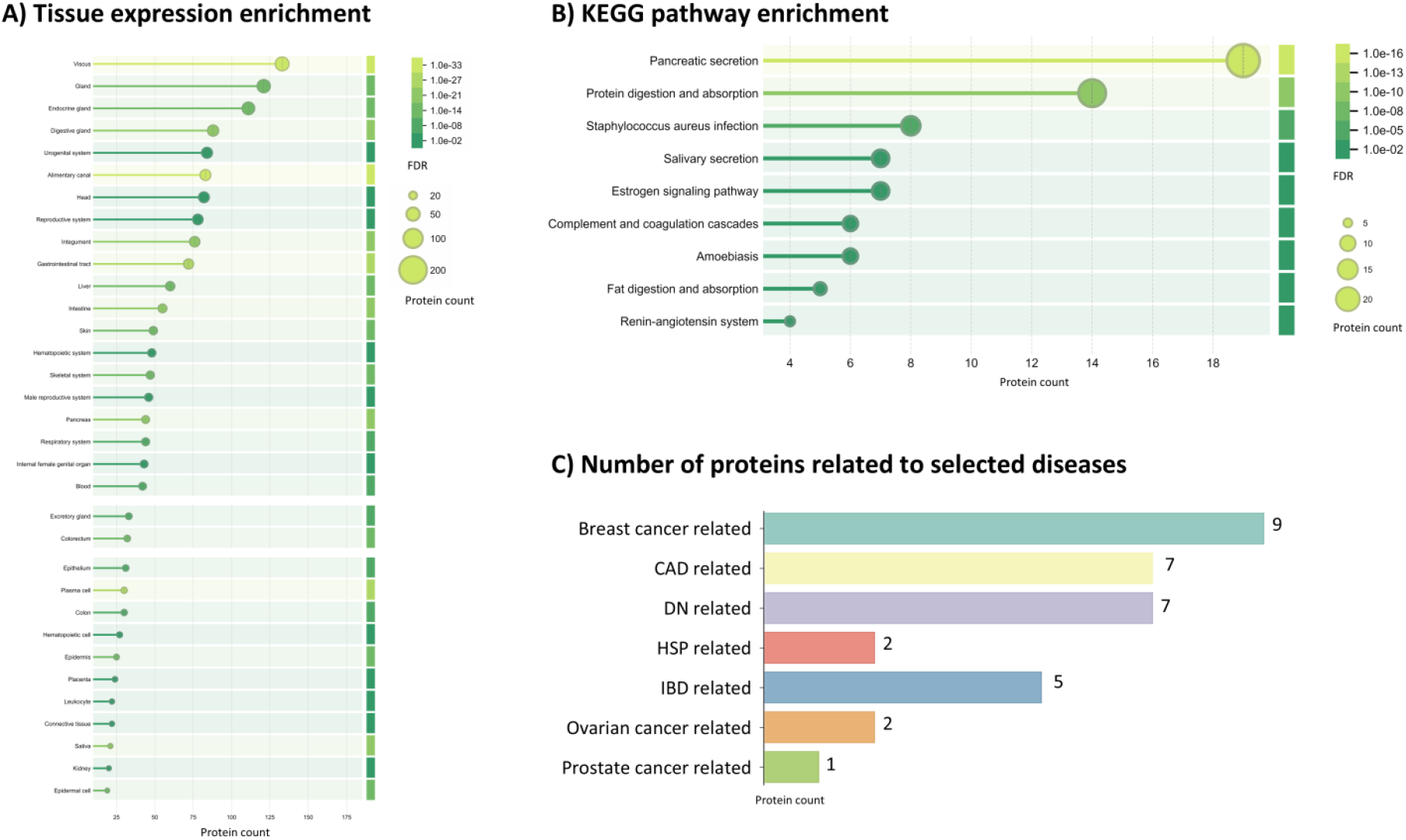
Analysis of the identified human proteins for enriched terms and functions. A) The graph displays the enrichment of tissue expression terms. The length of each bar and the size of the corresponding circle represent the number of proteins assigned to each term. The shade of green reflects the false discovery rate (FDR), as calculated by the STRING database tool. B) The graph illustrates the enriched KEGG pathways, with the bar length and circle size indicating the number of proteins assigned to each pathway. The shade of green represents the FDR, as determined by the STRING database tool. C) The bar graph shows the number of proteins associated with selected diseases. Abbreviations: CAD (Coronary Artery Disease), DN (Diabetic Nephropathy), HSP (Henoch-Schönlein Purpura), and IBD (Inflammatory Bowel Disease). The enrichment output tables as obtained from the STRING database tool are available in the SI EXCEL DOC.

Multiple proteins could be associated with breast cancer, followed by proteins linked to diabetic neuropathy, and some for inflammatory bowel disease (Figure 5C). Notable proteins with clinical relevance include arginase-1 (P05089), neutrophil defensins 1 (P59665), Calmodulin-like protein (Q9NZT1) and a range of immune response-related proteins. Potential health indicators are also the various immunoglobulin chains (e.g., from IgG, IgM, and IgA), which reflect inflammation or immune responses to pathogens, including viruses.

To gain a more comprehensive understanding of the enriched functions, tissue expression profiles, cellular locations, and pathways associated with the human proteins identified in wastewater, we conducted an enrichment analysis using the STRING database^54^ (Figure 5A and 5B, and SI EXCEL DOC). The analysis for tissue expression terms showed enrichments for gastrointestinal tract, digestive glands, skin, liver, pancreas, salivary glands, and respiratory system, reflecting their presence in bodily fluids such as saliva, urine, bile, and tears (Figure 5A). Notably, proteins were also linked to tissues involved in immune responses, such as bone marrow, blood, and plasma cells, along with specific tissues like epidermis, epithelial cells, and intestinal epithelium. Additionally, certain proteins were associated with reproductive tissues, including the prostate gland, ovary, and seminal plasma, as well as specific cell types like keratinocytes and leukocytes. This underscores the wide range of biological sources contributing to the human proteome detected in wastewater. Similarly, the enriched cellular component Gene Ontology reflects their extracellular and membrane-associated roles. Key terms include extracellular space, extracellular exosome, secretory granule, and vesicle, indicating that these proteins are actively secreted or involved in extracellular processes. Proteins were also linked to extracellular matrix, suggesting their role in tissue structure and integrity, and collagen-containing extracellular matrix, which may point to their involvement in connective tissues. Several terms related to granules, such as azurophil granule and zymogen granule, include protein storage and secretion functions.

Additionally, intermediate filament and keratin filament show that several of these proteins are involved in maintaining cellular architecture. Other terms like lysosome and autolysosome suggest potential involvement in protein degradation pathways. Enriched KO pathways include pancreatic secretion and protein digestion and absorption which are linked to the digestive system (Figure 5B). The identification of associations with *Staphylococcus aureus* infection and amoebiasis suggests the potential involvement of these proteins in immune responses to bacterial and parasitic infections. Salivary secretion and fat digestion and absorption show their involvement in digestive and metabolic functions. Furthermore, the renin-angiotensin system, complement and coagulation cascades, and estrogen signaling pathway were enriched, which link to cardiovascular regulation, immune responses, and hormonal signaling. The most prevalent molecular function Gene Ontology terms associated with the human proteins were as expected related to enzymatic and structural roles. Key terms included endopeptidase inhibitor activity, peptidase activity, and serine-type peptidase activity, indicating the involvement of these proteins in proteolytic processes. Several structural roles were also identified, such as structural constituent of skin epidermis and the cytoskeleton, proteins which maintain cellular integrity. Other relevant activities include metal ion binding, antioxidant activity, immunoglobulin binding, enzyme inhibitor activity, and toxic substance binding, suggesting diverse physiological roles for these proteins in health and disease processes.

Most interestingly, the human proteins detected in wastewater can be linked to a wide range of clinically relevant disease gene associations, underscoring their potential relevance for public health monitoring (Figure 5C). Several skin diseases associations were enriched, including conditions such as keratosis, bullous skin disease, and skin cancer. These proteins were also linked to respiratory system diseases, with lung disease and lower respiratory tract disease. The identified proteins could also be linked to diseases such as pancreatitis. Proteins related to various types of cancer are further outlined in SI Figure 2 and SI Table 1.

## Discussion and conclusions

The presented study demonstrates a streamlined wastewater metaproteomics approach which provides a comprehensive view of the wastewater metaproteome. This approach uses filter-aided sample preparation from small wastewater volumes and de novo sequencing to refine global reference sequence databases – which makes it suitable for multiplexing. We applied this approach to profile the wastewater metaproteome over three months from two different municipal locations. Interestingly, while clustering revealed distinct differences between individual time points and locations, a dominant core metaproteome remained consistent across all samples. This observation may be further amplified by the generic reference sequence database, which might not capture all differences. Additionally, such databases limit the exploration of microbial populations to the genus or family level. On the other hand, the core proteome is likely similar, as both municipal wastewater locations are in densely populated areas, only about 55 km apart, differing mainly in their sewer residence times. The identified proteins belong to environmental microbes as well as microbes form the human microbiome, such as the human gut. Among the many detected families, several contain pathogenic microbes and those known to spread antimicrobial resistance^53^. This may provide valuable insights into the spread of such genes in the environment. Since metaproteomics directly measures proteins, indicating that these microbes were either active or dormant, the information on the potential spread of antibiotic resistance genes would also be more relevant.

However, while metaproteomics identified a broad spectrum of microbes, the unambiguous identification of indicator strains or pathogens requires species-level resolution. This study used a public reference sequence database, which did not allow distinction between individual strains. Nevertheless, this could be achieved by utilizing metagenomic reference sequence databases in addition to the public UniRef database. Furthermore, the presented study did not include viruses, which are of great interest for controlling emerging pandemics. Viruses constitute only a tiny fraction of the protein biomass in such samples, and they typically produce only a few proteins. Consequently, a high viral load must be present in the wastewater for successful metaproteomic detection. Only a few studies employing targeted methods have reported the successful detection of SARS-CoV-2^92^. For example, Jagadeesan and co-workers demonstrated the simultaneous detection of SARS-CoV-2 proteins and the C-reactive protein, which next to the spread of the virus may also indicates inflammation responses^93^. Additionally, different antibody chains have been detected, such as the IgGFc-binding protein and the J chain from IgA, which links two monomer units of either IgM or IgA, and which may also indicate the presence of a viral infection in the population. Stephenson and colleagues explored the detection of antibodies in wastewater and evaluated their immunoaffinity against the SARS-CoV-2 spike protein^94^. They observed intact antibodies, predominantly of the IgG and IgA classes, which appeared to adhere to solid matter in the wastewater. IgA antibodies, critical for mucosal immunity, are detectable in the respiratory tract and serum within one to two weeks following infection or vaccination. In contrast, IgG antibodies typically emerge between two to three weeks and persist for several months. The study demonstrated that these IgG and IgA antibodies retained their immunoaffinity for SARS-CoV-2 spike protein antigens^94^. This suggests that active antibody fractions in wastewater could facilitate real-time monitoring of population immunity, vaccination coverage, and infection prevalence. This may help assess the effectiveness of vaccination campaigns and complement traditional seroprevalence surveys, which are time-intensive to conduct.

The detection of a wide spectrum of human proteins is another important aspect of wastewater metaproteomics. This data may support the evaluation of population health or signal the emergence of an epidemic. In this study, we identified approximately 200 human protein groups, including several potential disease-related proteins and biomarkers, which are promising indicators of population health. Recent studies have shown that population averages for different protein biomarkers can be estimated with good accuracy for several diseases^95 96^. However, projecting the detected wastewater concentrations to individual averages requires consideration of dilution rates and the population size in the area. It has been suggested that using an abundant and easily detectable human protein, such as albumin, as a normalization factor could further improve protein quantification in wastewater^50^. It is worth mentioning that, in the case of the human proteome, diseases may also correlate with changes in protein modifications. For example, alterations in protein glycosylation have been extensively studied for their relevance as biomarkers^97 98 99^. However, no studies to date have investigated the fate of these modifications in the complex wastewater environment.

Besides the advantage of anonymity, wastewater collection is relatively inexpensive and can be applied to large population sizes. However, wastewater collected from extensive areas, including both urban and rural regions, tends to have a longer retention time in the pipes and likely a more uniform composition. For the implementation of wastewater surveillance, city-proximal sampling points may also allow to observe subtle changes in less prevalent pathogens or protein biomarkers. Finally, although mass spectrometry-based metaproteomics is a powerful approach for both fully untargeted screening but also sensitive targeting, this approach still shows some limitations towards throughput and sensitivity. Successful detection requires a minimum number of protein copies, making low-abundance microbial, viral and human proteins challenging to detect. At the same time, the developments in mass spectrometric instrumentation and methods has significantly improved sensitivity and throughput, which is highly relevant when monitoring complex and dynamic environments such as wastewater^42, 100, 101^. Furthermore, alternative technologies are currently being developed. For example, significant advancements have been made in single-protein sequencing technologies, such as the use of nanopores, similar to DNA sequencing^102^. Progress has also been made in the development of single-protein fingerprinting approaches, which additionally may provide insights into protein modifications^103^. These advancements hold great promise for the integration of metaproteomics into routine wastewater monitoring in the near future.

## Supporting information

SI WORD DOC

SI EXCEL DOC

## Acknowledgements

The authors thank Dita Heikens for her support in the proteomics laboratory and acknowledge the NWO Spinoza Prize awarded to Mark van Loosdrecht for funding. They also thank Jelle Langedijk and Mario Pronk for their assistance in obtaining wastewater and extend special thanks to the engineers at the Utrecht and Harnaschpolder wastewater treatment plants for their help with sampling.

