## Supplementary material for "Wastewater metaproteomics: tracking microbial and human protein biomarkers": SI WORD DOC

### Supplementary information material to:
### Wastewater metaproteomics: tracking microbial and human protein biomarkers

Claudia G. Tugui, Filine Cordesius, Willem van Holthe, Mark C.M. van Loosdrecht and Martin Pabst*
Department of Biotechnology, Delft University of Technology, 2629 HZ Delft, The Netherlands.
*Contact:

**TABLE OF CONTENTS**

SI Figure 1A: LDA analysis of potential pathogens Page 2
SI Figure 1B: LDA analysis of human proteins Page 2
SI Figure 2: Proteins in HP and UT related to different types of cancers Page 3
SI Figure 3A: PCA analysis of human proteins identified in HP Page 4
SI Figure 3B: PCA analysis of human proteins identified in UT Page 4
SI Figure 4A: PCA analysis of metaproteome (excluding human proteins) identified in HP Page 5
SI Figure 4B: PCA analysis of metaproteome (excluding human proteins) identified in UT Page 5
SI Table 1: Proteins associated with different types of cancers in HP and UT Page 6

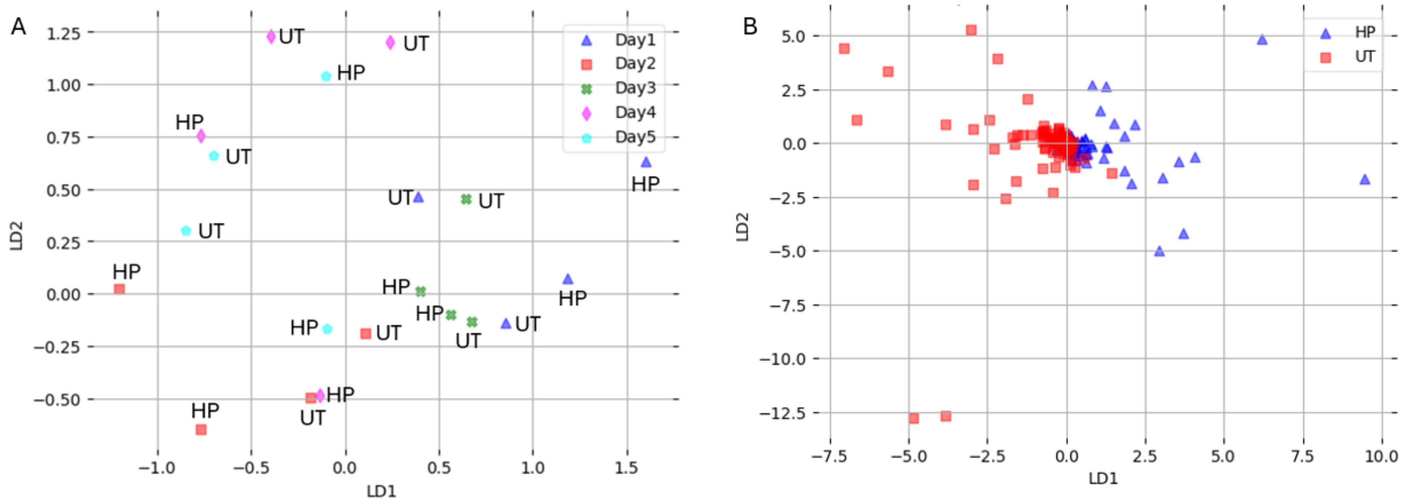


**SI Figure 1. A)** LDA analysis on potential pathogens (as defined by WHO report) discovered in wastewater samples based on sampling days. **B)** LDA analysis on the detected human proteome based on location (UT = Utrecht, HP = Harnaschpolder).

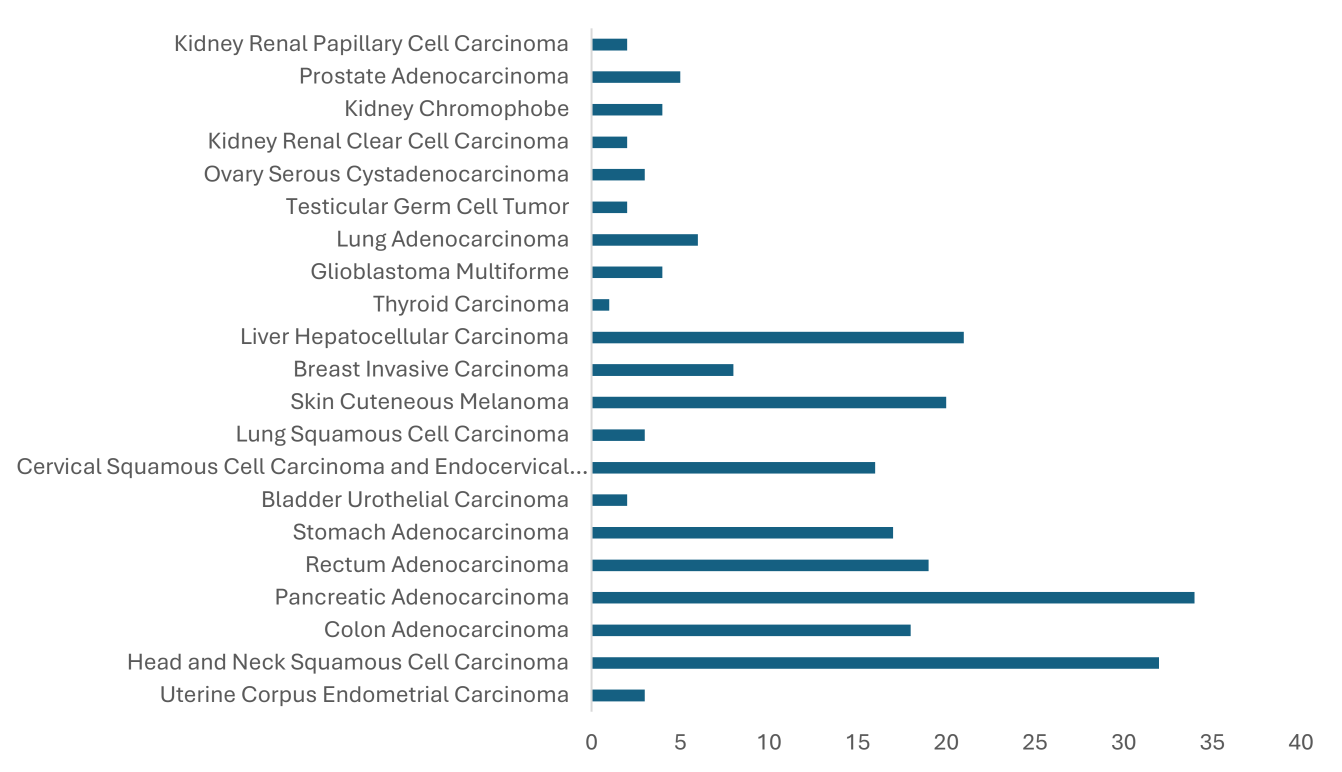


**SI Figure 2.** Number of human proteins with potential clinical relevance (e.g. biomarkers) for different types of cancer by both locations.

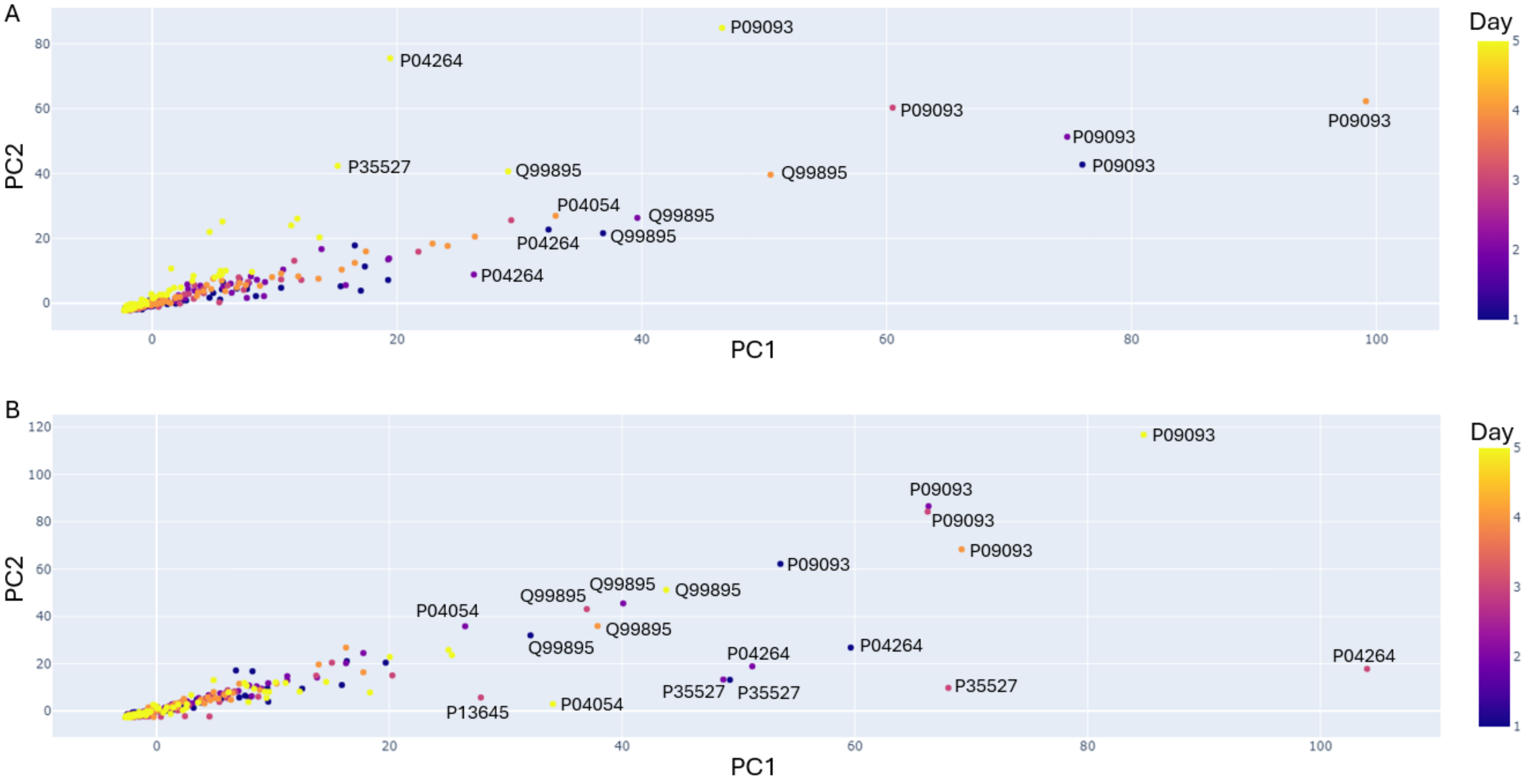


**SI Figure 3.** PCA analysis of the detected human proteome for HP (Graph A) and UT (Graph B). Outliers are annotated by their UniProtKB accession number. The color of the dot represents the day of sampling.

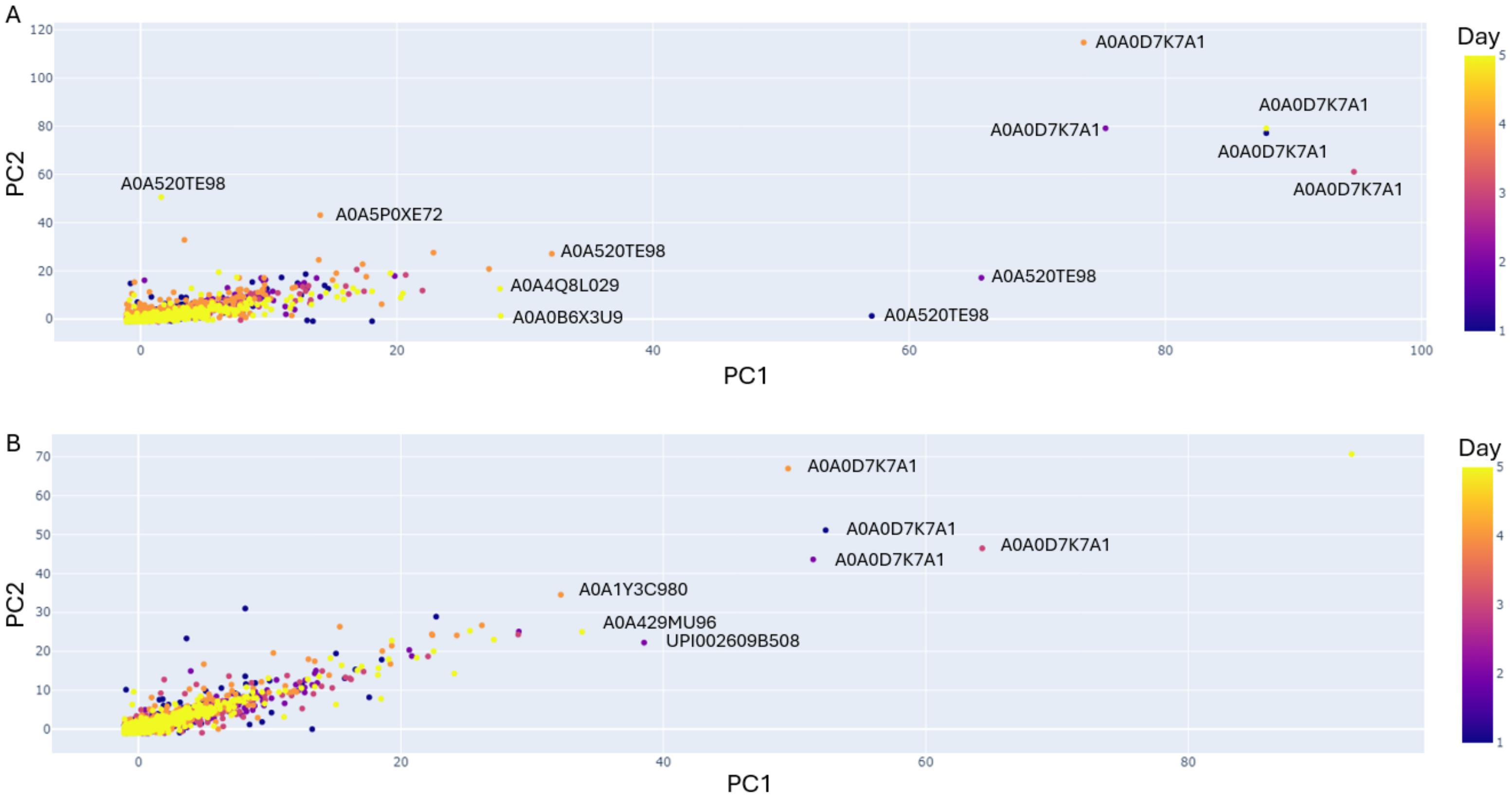


**SI Figure 4.** PCA analysis of the detected microbial metaproteome for HP (Graph A) and UT (Graph B). Outliers are annotated by their UniProtKB accession number. The color of the dot represents the day of sampling.

**SI Table 1.** List of proteins potentially related to different types of cancer, as classified by the Human Protein Atlas (UT = Utrecht, HP = Harnaschpolder).

| Cancer type | UniProtKB accessions of potentially related proteins |
|---|---|
| Uterine Corpus Endometrial Carcinoma | Q14508, P01036, P20160 |
| Head and Neck Squamous Cell Carcinoma | P04264, P15924, P13645, P08779, P02533, P04259, P13647, Q04695, Q02413, P13646, P29508, Q8N1N4, P06702, Q01546, Q5T749, Q08554, Q08188, Q9NZT1, P04083, Q01469, P31944, P31151, Q96P63, P05109, P35609, Q13835, P22532, A8K2U0, P01040, Q15517, P63316, P55000 |
| Colon Adenocarcinoma | Q9Y6R7, Q02817, A8K7I4, Q9UGM3, Q16819, Q8WWA0, P35030, P56470, O60844, P06731, P40879, P05451, P00915, Q14002, P13688, Q12864, Q9H3R2, Q6UX06 |
| Pancreatic Adenocarcinoma | Q02817, P04746, P19961, A8K7I4, Q9UGM3, Q86UP6, P08861, P09093, P07477, P55259, P16233, P07478, Q6GPI1, P08217, P15085, P98088, Q8WWA0, Q99895, P48052, P04054, P98073, Q16820, P35030, P56470, P05451, Q6W4X9, P04118, Q03403, P19835, P02766, Q9H3R2, P31025, Q9HD89, Q6UX06 |
| Rectum Adenocarcinoma | Q02817, A8K7I4, Q9UGM3, Q16819, Q8WWA0, Q8WWU7, P35030, P56470, O60844, P06731, P40879, P00915, Q14002, P13688, Q12864, P22748, P40199, Q9H3R2, Q6UX06 |
| Stomach Adenocarcinoma | Q02817, Q9UGM3, P14410, P98088, Q8WWA0, P05164, P98073, P35030, P61626, P56470, P05451, P80188, Q12864, Q6W4X9, Q03403, Q9H3R2, Q6UX06 |
| Bladder Urothelial Carcinoma | P13646, P31944 |
| Cervical Squamous Cell Carcinoma and Endocervical Adenocarcinoma | P13647, Q04695, Q14CN2, P13646, P29508, P48594, Q8N1N4, P06702, Q9NZT1, Q01469, P31944, Q96P63, P05109, Q13835, P01040, Q9HCY8 |
| Lung Squamous Cell Carcinoma | P13646, Q96P63, Q13835 |
| Skin Cuteneous Melanoma | P04264, P35527, P35908, P13645, P04259, Q86YZ3, Q02413, Q5D862, Q8N1N4, P05090, Q5T749, Q08554, Q9NZT1, P31944, P31151, Q96P63, O14556, Q15517, P20930, P55000 |
| Breast Invasive Carcinoma | P02788, P25311, P12273, Q96DA0, P00709, P01036, P81605, P02814 |
| Liver Hepatocellular Carcinoma | P02768, P02787, P01009, P09923, O43895, P25311, Q6UWV6, P01011, P02760, P01008, P00738, P56470, P05089, P02763, O95497, P05154, P00734, P02790, P02766, P02750, P05155 |
| Thyroid Carcinoma | P61916 |
| Glioblastoma Multiforme | P01011, P35609, P10153, P59665 |
| Lung Adenocarcinoma | P0DTE8, Q9UGM3, P98088, Q8WWA0, P40199, Q9HD89 |
| Testicular Germ Cell Tumor | Q8WWU7, Q6PEY2 |
| Ovary Serous Cystadenocarcinoma | P0DTE8, Q14508, P31025 |
| Kidney Renal Clear Cell Carcinoma | P09923, P07911 |
| Kidney Chromophobe | P07911, P06870, P24855, P01133 |
| Prostate Adenocarcinoma | P27487, P25311, P15309, Q96DA0, P14555 |
| Kidney Renal Papillary Cell Carcinoma | P15144, Q9BYE9 |
